# ENAH Binding to the Huntingtin Proline-Rich Domain Modulates HTT1a Assembly

**DOI:** 10.64898/2026.07.29.741051

**Authors:** Jobin Varkey, Anoop Rawat, Priyatama Pandey, Baiyi Quan, Tsui-Fen Chou, Ralf Langen, Ali Khoshnan

**Author notes:** For correspondence: Ali Khoshnan.

## Abstract

The proline-rich domain (PRD) of huntingtin (HTT), located C-terminal to the polyglutamine tract within the exon-1 region, plays a critical role in modulating the aggregation and toxicity of mutant HTT1a. To identify PRD-interacting proteins, we performed mass spectrometry analysis of PRD peptide pulldowns from human neurons. ENAH, a member of the Ena/VASP family of actin regulatory proteins, emerged as the top enriched interactor from neuronal membrane fractions. In neurons, ENAH colocalized with HTT1a species, suggesting a biologically relevant interaction. Because the EVH1 domain of ENAH binds polyproline motifs with high affinity, we examined its interaction with HTT1a using recombinant proteins. EVH1 directly bound HTT1a and unexpectedly formed condensate-like assemblies capable of recruiting HTT1a monomers. Remarkably, EVH1 robustly suppressed HTT1a oligomerization and promoted remodeling of preformed HTT1a fibrils in vitro. These findings demonstrate that ENAH–PRD interactions modulate HTT1a aggregation dynamics and identify EVH1 assembly as a potential proteostatic mechanism regulating mutant HTT1a states. More broadly, this work establishes a framework for investigating how EVH1-containing proteins may influence HTT1a proteostasis and neurotoxicity in Huntington′s disease.

## INTRODUCTION

Huntington’s disease (HD) is caused by the expansion of the CAG repeat in exon-1 of the huntingtin gene, producing a protein with an expanded polyglutamine (polyQ) repeat (*1*). Huntingtin exon-1 fragment (HTT1a) is predominantly translated from an aberrantly spliced HTT mRNA and accumulates as aggregates in the brains of HD patients (2–4). Recent work further suggests that premature or abortive translation termination of expanded HTT transcripts can generate truncated, aggregation-prone N-terminal HTT fragments, providing an additional source of mutant HTT species (5). HTT1a contains an N-terminal amphipathic segment (N17), a polyglutamine (polyQ) tract, and a proline-rich domain (PRD) at the C-terminal end. HTT1a with expanded polyQ has increased aggregation propensity, resulting in the formation of HTT1a fibrils (*6*). Structural models show that the fibrillar HTT1a consists of a core of tightly packed β-sheets formed by the polyQ region, with the C-terminal PRD extending outwards like bristles (*6–8*). HTT1a species are known to sequester cellular proteins, likely including proteins trapped by binding to PRD bristles, and influence the oligomerization and toxicity of HTT1a (*9–11*).

To identify endogenous PRD binders of HTT1a in a neuronally relevant context, we performed a pulldown–mass spectrometry screen with HTT1a PRD as bait using multiple fractions of human neuronal lysates. This unbiased proteomic survey identified ENAH, also known as mammalian Enabled/MENA, as a significantly enriched candidate across multiple fractions and as the top interactor in neuronal membrane fractions. ENAH is a member of the Enabled/vasodilator-stimulated phosphoprotein (Ena/VASP) family of actin regulatory proteins. Vertebrate Ena/VASP proteins regulate actin assembly and cytoskeletal remodeling during cell migration, neuronal development, axon outgrowth, and synaptic organization (12–15). Ena/VASP proteins are recruited to different regions of the cell by binding proline-rich SLiMs (Short linear motifs) via their N- terminal EVH1 domains (*16, 17*). EVH1 domains from the Ena/VASP family canonically recognize FP4-like motifs but can accommodate extended and noncanonical sequences in multivalent contexts such as the PRD domain of HTT1a (16,17).

Here, employing recombinant proteins, we examined the interaction between the EVH1 domain of ENAH and HTT1a and assessed its potential influence on HTT1a assembly behavior. Our findings suggest that EVH1-mediated engagement of the PRD may constitute an additional layer of regulation in HTT1a proteostasis and raise the possibility that ENAH condensation may contribute to this process.

## MATERIALS AND METHODS

### PRD pulldown and mass spectrometry

A recombinant fragment corresponding to the huntingtin proline-rich domain (PRD) of HTT1a was expressed in *E. coli* with an N-terminal His₆ tag, purified by nickel affinity chromatography, and immobilized on Ni²⁺-NTA magnetic beads. Bead-bound His-PRD was incubated with the membrane fractions prepared from cultured human neurons under native binding conditions (*18*). After extensive washing to remove nonspecific proteins, bound complexes were eluted and subjected to tryptic digestion followed by LC–MS/MS analysis. To control for nonspecific interactions, parallel pulldowns were performed using beads loaded with an unrelated His₆-tagged IL-34. Proteins uniquely enriched in the His-PRD pulldown relative to control were considered specific PRD-binding candidates.

### Quantification of proteomic data

R version 4.4.2 was used for coding and analysis of quantitative proteomic data. Differentially enriched proteins were identified using an adjusted p-value cutoff of <0.05 and a fold-change threshold of 1.5. Volcano plots were generated using the Enhanced Volcano package (version 1.24.0) to visualize significant changes and highlight the top differentially enriched proteins. Volcano plots were generated for all five datasets, applying the same statistical cutoffs (adjusted p-value <0.05, fold change ≥1.5). In addition, for each dataset, an Excel file containing three sheets was prepared: (1) “de_output_adj_p_value_0.05” for all differentially enriched proteins, (2) “upregulated_adj_p_value_0.05” for upregulated proteins, and (3) “downregulated_adj_p_value_0.05” for downregulated proteins.

#### Protein expression and purification

The human ENAH EVH1 domain with the coiled-coil sequence, which was previously cloned into a vector, pDW363, was obtained from Prof. Amy Keating’s lab (MIT, Boston) (*16, 17*). The expressed protein is labeled as EVH1. This construct was transformed into Rosetta2(DE3) (Novagen) cells, and the transformed cells were grown overnight in Terrific Broth with 100 μg/mL ampicillin at 37 °C. Then the cells were expanded at 37 °C and induced with 0.5 mM IPTG at an optical density of ∼0.8 at 600 nm. The expression was carried out at 37 °C for 5 h. The cells were then spun down and resuspended in lysis buffer consisting of 20 mM Hepes pH 7.4, 500 mM NaCl, 20 mM imidazole, 2 mM DTT, and frozen at –80 °C overnight for long-term storage. The next day, pellets were thawed followed by addition of 100 mM phenylmethylsulfonyl fluoride (Sigma) and protease inhibitor tablets (Pierce). Bacterial cells were lysed by using a tip sonicator and the lysate was centrifuged at 19000 g for 20 min. The supernatant was incubated with NiHis60 (Takara Bio) for 1 h. The protein was eluted using 50 mM Hepes pH 7.4, 50 mM NaCl, 500 mM imidazole, 2 mM DTT. The eluate was dialyzed into 20 mM Hepes pH 7.4, 50 mM NaCl, 2 mM DTT buffer. The protein was further purified using Superdex 75 10/300 gel filtration column using 20 mM Hepes pH 7.4, 50 mM NaCl, 2 mM DTT buffer.

### Dot blot binding assay

For dot blot analysis, equal molar amounts of monomeric or fibrillar HTT1a (46Qs) were diluted in PBS and spotted (1–2 µL per spot) onto nitrocellulose membranes. Membranes were air-dried for 15 minutes and blocked for 1 hour at room temperature in 5% nonfat dry milk in TBST (Tris-buffered saline with 0.1% Tween-20) (*6, 19*). Blocked membranes were incubated overnight at 4 °C with recombinant EVH1 domain of human ENAH (0.5–1 µM) diluted in blocking buffer to allow binding. After washing three times with TBST, membranes were probed with anti-ENAH or anti-His antibody (when His-tagged EVH1 was used), followed by HRP-conjugated secondary antibody for 1 hour at room temperature. Signal was detected using enhanced chemiluminescence.

### Cell culture

Human MESC2.10 neural progenitor cells (NPCs) were maintained under proliferative conditions in NPC expansion medium consisting of DMEM/F12 supplemented with N2, B27 (minus vitamin A), GlutaMAX, penicillin/streptomycin, and basic fibroblast growth factor (bFGF). Cells were cultured at 37 °C in a humidified 5% CO₂ incubator and passaged every 3–4 days. For neuronal differentiation, NPCs were plated onto poly-D-lysine/laminin–coated coverslips and cultured in differentiation medium lacking bFGF. Cells were differentiated for 7 days prior to analysis (18–20).

### Lentiviral vector construction and production

HTT1a with 72Q length or human ENAH cDNA were cloned into a lentiviral backbone as described previously (*18–20*). Where indicated, ENAH was fused at the C-terminus to a fluorescent reporter (or EGFP) via a flexible (GGGGS)₃ linker to allow visualization without disrupting EVH1 function. Replication-deficient lentivirus was produced by co-transfecting HEK293T packaging cells as described and titrated (*18–20*).

### Lentiviral transduction of NPCs

MESC2.10 NPCs were transduced with lentivirus encoding HTT1a-72Q. Similarly, NPCs with or without HTT1a were transduced with lentiviruses encoding ENAH or EGFP-ENAH at a multiplicity of infection (MOI) of 2:1 in the presence of 8 µg/mL polybrene, to ensure high transduction. After 24 hours, medium was replaced with fresh expansion medium. Cells were allowed to recover for 48 hours before initiating neuronal differentiation. Expression of ENAH was confirmed by immunofluorescence using antibodies against ENAH or the fluorescent tag. HTT1a expression was confirmed by antibody staining using PHP1 (*19*).

### Immunofluorescence and confocal microscopy

Differentiated neurons were fixed in 4% paraformaldehyde for 15 minutes at room temperature, permeabilized with 0.1% Triton X-100, and blocked with 5% normal goat serum. Cells were incubated overnight at 4 °C with primary antibodies against HTT1a (18), and ENAH (ThermoFisher, HA500026, Waltham, MA). After washing, appropriate Alexa Fluor–conjugated secondary antibodies were applied for 1 hour at room temperature. Coverslips were mounted using antifade reagent containing DAPI. Images were acquired using a laser-scanning confocal microscope with a 60× oil immersion objective. Z-stacks were collected to assess cytoplasmic and neuritic localization. EGFP-ENAH transduced cells were visualized live by fluorescent microscope.

#### Fluorescence microscopy

Concentrated stocks of EVH1 were diluted into the required concentrations in 20 mM Hepes, 50 mM NaCl, pH 7.4, with 10% PEG 4000. Samples were then imaged using a Zeiss Axioplan fluorescence microscope at 40x. In case of co-mixing experiments, the HTT1a(Q25) monomers were labeled with Alexa-488.

#### Transmission electron micrography

The samples were diluted tenfold and applied onto carbon-coated copper electron microscopy (EM) grids and stained using 1% uranyl acetate. Negative staining images were captured using a JEOL JEM-1400 transmission electron microscope at 100 kV.

#### Thioflavin T fluorescence assay

We used Thioflavin T at a 50 μM final concentration to monitor HTT1a-Q46 misfolding (*5*). HTT1a-Q46 was used at a concentration of 10 μM while EVH1 was used at 20 μM. The aggregation kinetics were performed in Nunc black 96-well optical bottom plates (Thermo Scientific) in FLUOstar OMEGA, BMG Labtech microplate reader, with measurements every 15 minutes. All assays were carried out at 25 °C in 20 mM TBS buffer at a final well volume of 120 μl.

## RESULTS

### Identification of PRD interactors

Mass spectrometry analysis of proteins recovered from His-PRD pulldowns revealed multiple candidates enriched significantly over control conditions (Fig. 1). Among these, ENAH, an actin-regulatory protein of the Ena/VASP family, was identified as an interactor in multiple organelles and was the top candidate in the neuronal membrane fractions, suggesting a preferential association with the HTT1a PRD. Vasodilator-stimulated phosphoprotein (VASP), another key actin-regulatory protein of the Ena/VASP family (12), was also significantly enriched in the PRD pulldown along with previously identified PRD binding protein profilin (Supplementary File 1). These results identify ENAH, a major regulator of cytoskeletal dynamics, and potentially other Ena/VASP family members as new interactors of the PRD in HTT.

**Figure 1.**
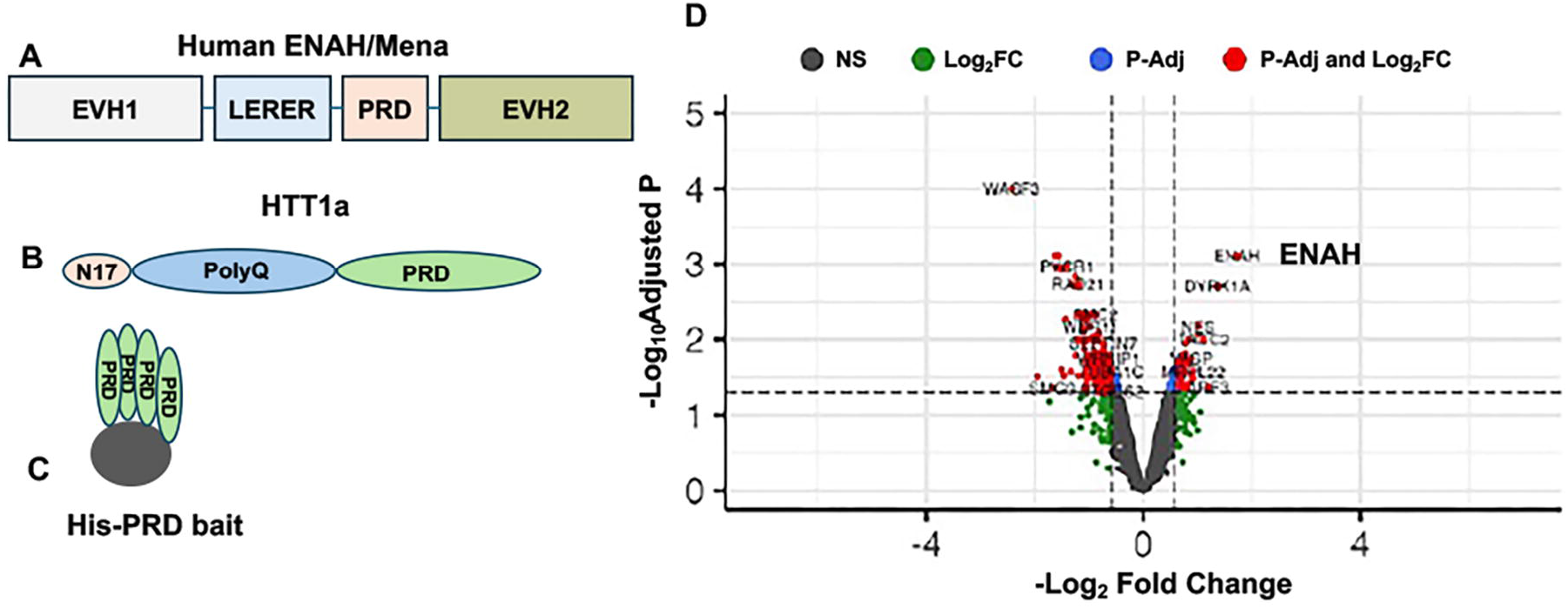
PRD pulldown of neuronal lysates identifies ENAH as an enriched HTT1a PRD interactor. (A) Schematic representation of human ENAH/Mena domain organization. (B) Schematic representation of HTT1a, showing the N17, polyQ, and PRD regions. (C) Schematic of PRD immobilized on magnetic beads for pulldown of PRD-interacting proteins. (D) Volcano plot of mass spectrometry data from PRD pulldowns showing enriched candidate interactors, including ENAH.

### HTT1a colocalizes with ENAH in human neuronal cells

To examine the spatial relationship between HTT1a and ENAH in a neuronal context, we engineered a human stem cell–derived neuronal progenitor cell line (MESC2.10) to express HTT1a containing 72 polyglutamine repeats together with human ENAH using lentiviral transduction (18–20). Immunofluorescence revealed substantial colocalization of HTT1a and ENAH in the cytoplasm and along neuritic extensions of differentiating neurons (Fig. 2). Interestingly,

**Figure 2.**
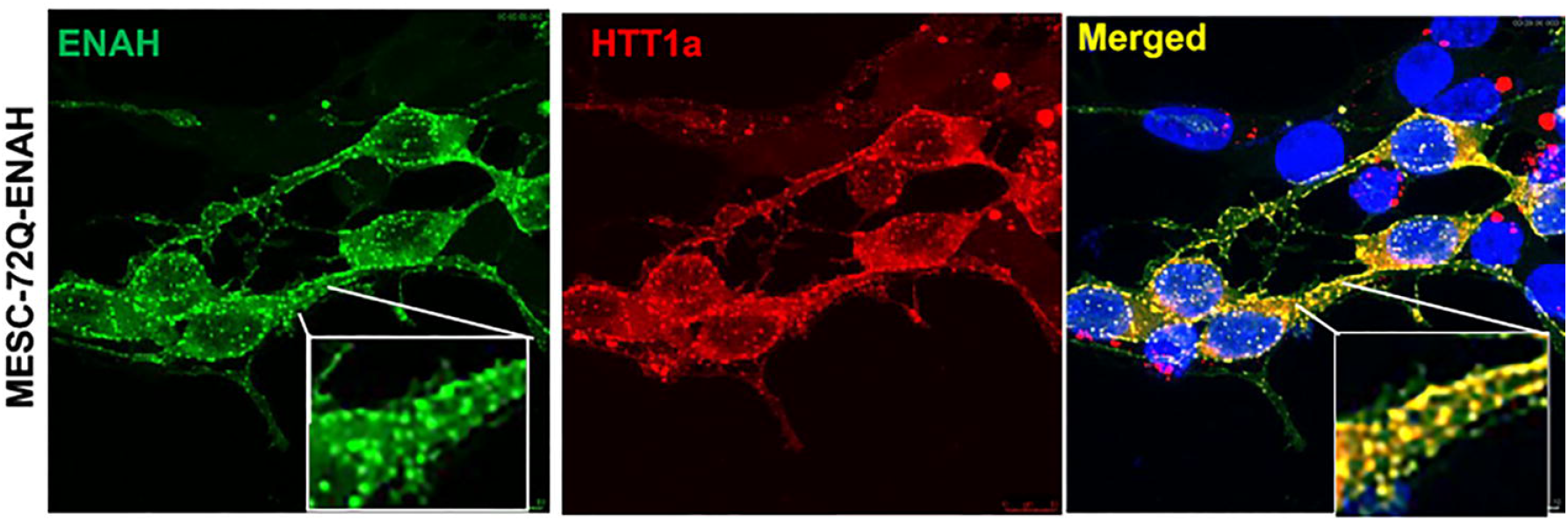
ENAH colocalizes with HTT1a species in human neurons. MESC2.10 neuronal progenitor cells (NPCs) were engineered to express HTT1a (72Q) together with human ENAH using lentiviral transduction and differentiated into neurons. At day 7 of differentiation, neurons were immunostained with rabbit anti-ENAH or anti-HTT1a (PHP1) antibodies and imaged by confocal microscopy. Insets, magnified regions showing ENAH condensate-like assemblies (left panel) and colocalization with HTT1a (merged).

ENAH appeared as small, distinct puncta characteristic of higher-order assemblies (Fig. 2 inset). To determine whether ENAH forms these structures independently of HTT1a in live cells, we expressed EGFP-ENAH in the presence or absence of HTT1a and performed live-cell imaging. We found that ENAH forms puncta-like structures under both conditions (Fig. 3). These observations indicate that ENAH can assemble into higher-order structures independently of HTT1a, while HTT1a can localize to ENAH-enriched puncta in neurons.

**Figure 3.**
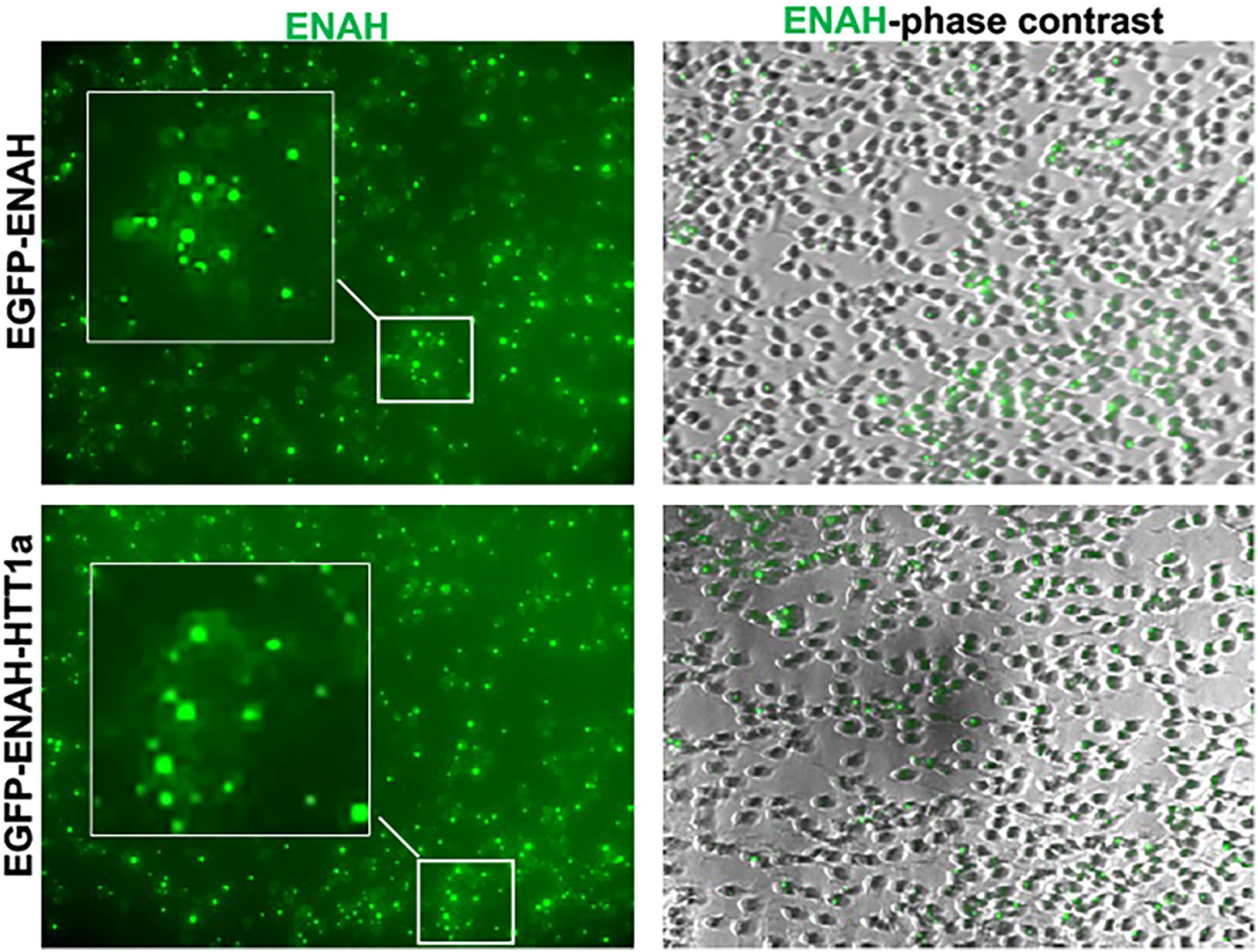
ENAH forms puncta-like structures independently of HTT1a. MESC2.10 neuronal progenitor cells were transduced with EGFP–ENAH in the presence or absence of HTT1a (72Q). Live-cell imaging revealed that EGFP–ENAH forms puncta-like structures under both conditions. Insets, magnified regions highlighting ENAH condensate-like assemblies.

### EVH1 binds to HTT1a monomers and fibrils

To determine whether EVH1 recognizes distinct HTT1a assembly states, we performed dot blot analysis using recombinant His-tagged EVH1 as probe. At equimolar concentrations, EVH1 showed binding to HTT1a monomers and fibrils (Fig. 4). The apparent increased binding to fibrils may reflect multivalent engagement of EVH1 with PRD motifs, which are repetitively displayed and accessible on the surface of HTT1a fibrils (7, 21).

**Fig. 4.**
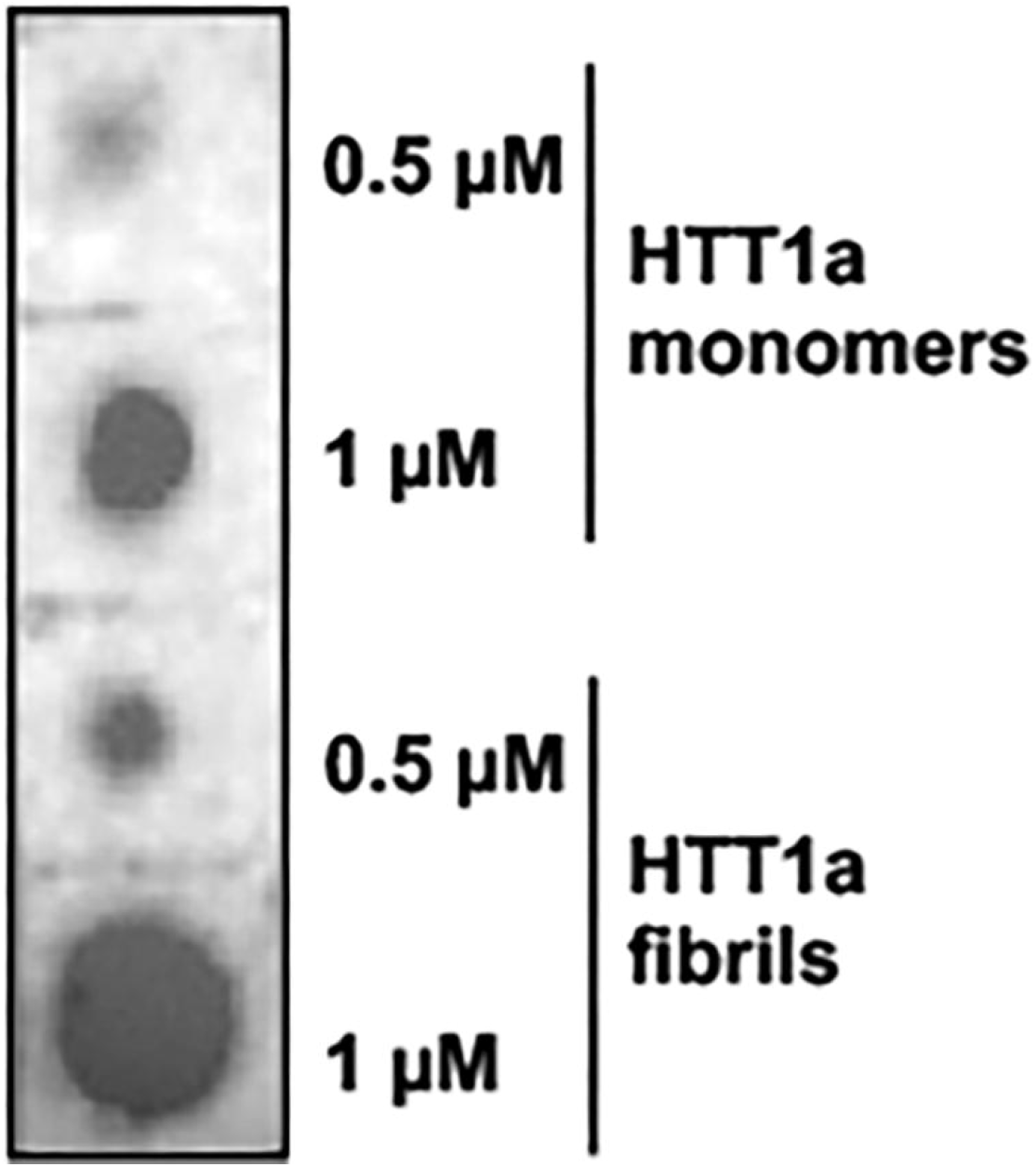
Dot blot analysis of EVH1 binding to HTT1a species. Serial amounts of monomeric and fibrillar HTT1a-Q46 were spotted onto nitrocellulose membranes and incubated with recombinant human EVH1. Bound EVH1 was detected using anti-His antibody followed by HRP-conjugated secondary antibody and chemiluminescence.

#### EVH1 forms condensates and recruits HTT1a monomers

The Ena/VASP family proteins, and other mammalian proteins such as Homer, contain an Enabled/VASP homology 1 (EVH1) domain that mediates direct binding to proline-rich motifs (16,17). Because ENAH formed punctate, condensate-like structures in human neurons (Figs. 2, 3), we asked whether recombinant EVH1 could form similar assemblies in vitro. Under macromolecular crowding conditions (22), EVH1 formed condensates (Fig. 5 A, B). Importantly, the EVH1 condensates recruited soluble HTT1a monomers (Fig. 5 C, D). This condensate-mediated recruitment of HTT1a suggests that EVH1 may act as a dynamic scaffold that concentrates mutant HTT1a in compartments, thereby facilitating its sequestration.

**Figure 5.**
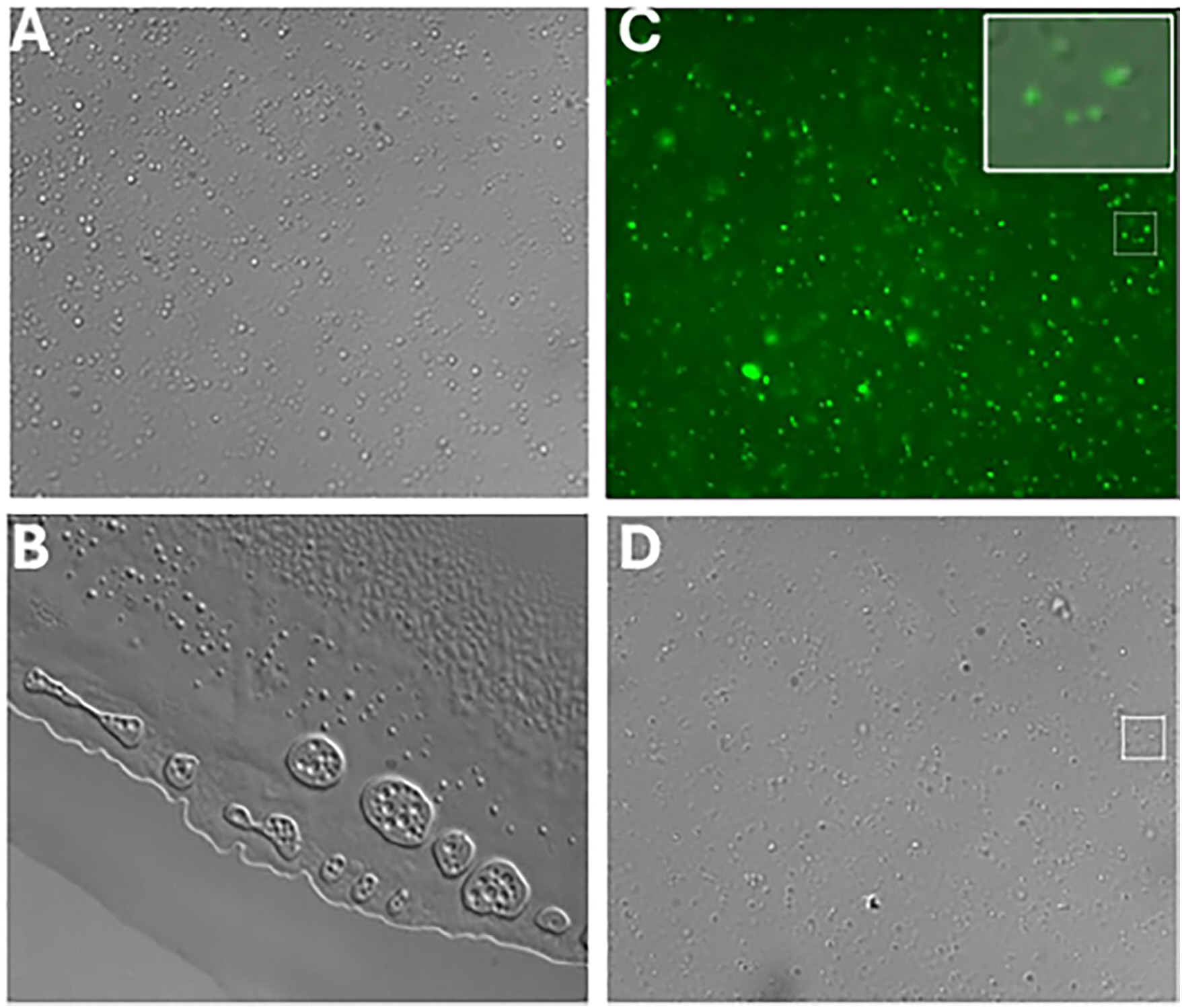
EVH1 forms condensates. (**A**) Formation of condensates. (B) Gel-like condensates after 30 min. (C, D) HTT1a-Q25 monomers partition into EVH1 condensates. Panel C is a fluorescence image, while panel D is a DIC image. Inset, magnified region showing monomeric HTT1a associated with EVH1 condensate-like assemblies.

### EVH1 prevents aggregation of HTT1a-Q46

To examine whether EVH1 binding to PRD of HTT1a affects its aggregation, we performed Thioflavin T aggregation assays. We found that EVH1 significantly reduced the aggregation of HTT1a (Q46), while EVH1 alone did not exhibit any intrinsic aggregation (Fig. 6A). This inhibitory effect was further confirmed by TEM, which revealed abundant fibrils with HTT1a alone but no detectable aggregates in the presence of EVH1 (Fig. 6B-E). These findings highlight a potential role for EVH1-containing proteins binding to PRD on the assembly state of HTT1a.

**Figure 6.**
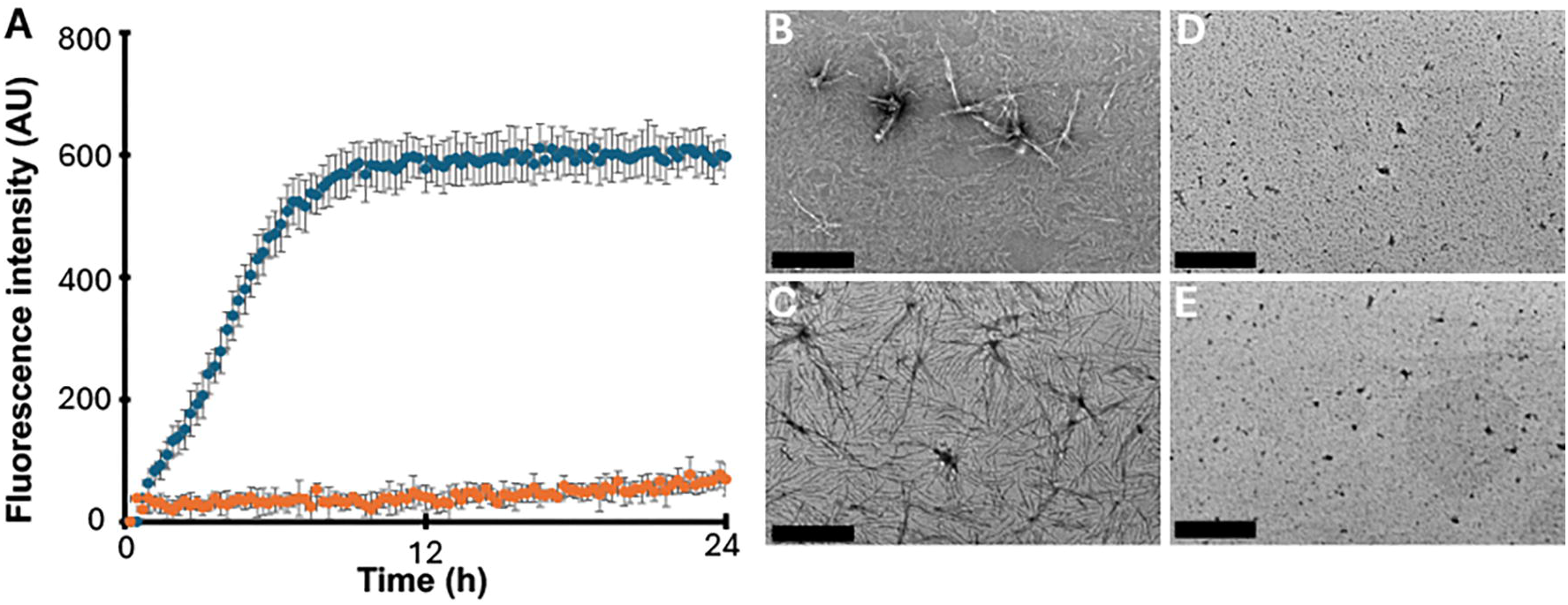
EVH1 inhibits HTT1a aggregation. (A) HTT1a-Q46 aggregation monitored using thioflavin T. The aggregation of 10 μM HTT1a alone (blue markers) was compared to that in the presence of 20 μM EVH1 (orange markers). Transmission electron micrographs were obtained for 10 μM HTT1a alone after 1 h (B), and 18 h (C), and compared to 10 μM HTT1a in the presence of 20 μM EVH1 after 1 h (D) and 18 h (E). The scale bar in B-D is 500 nm.

#### EVH1 can remodel preformed HTT1a-Q46 fibrils

Since EVH1 showed binding to HTT1a fibrils (Fig. 4), we examined the potential consequence of these interactions. When mixed at a 1:1 molar ratio, EVH1 promoted rapid remodeling and unbundling of HTT1a fibrils, which was evident within 10 minutes (Fig. 7A–C). Prolonged incubation for 15 hours resulted in extensive fibril remodeling, with marked unbundling and loss of fibrillar integrity (Fig. 7B–D). These findings suggest that EVH1–PRD interaction promotes fibril remodeling and unbundling, potentially shifting HTT1a assemblies toward non-fibrillar or more dispersed states.

**Figure 7.**
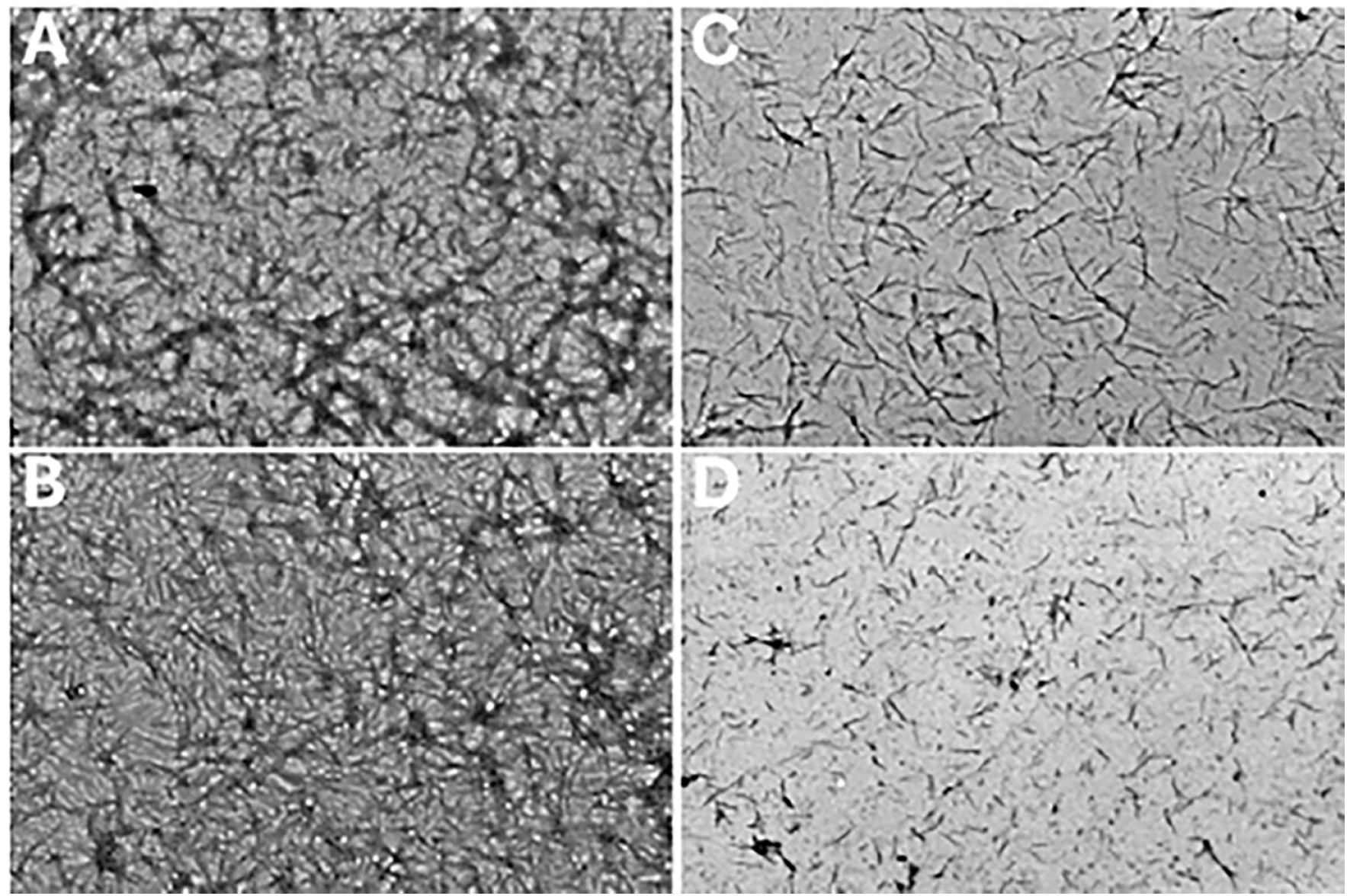
EVH1 unbundles preformed HTT1a fibrils. Transmission electron micrographs were obtained for 10 μM HTT1a preformed fibrils alone after 10 min (A), and 15 h (B), and compared with 10 μM HTT1a fibrils in the presence of 20 μM EVH1 after 10 min (C) and 15 h (D). The scale bar is 500 nm.

## DISCUSSION

In this study, we identified the Ena/VASP family protein ENAH as a prominent interactor of the huntingtin PRD and uncovered an unexpected role for its EVH1 domain in modulating the assembly state of HTT1a. Together, our biochemical and cellular data support a model in which EVH1 binding and potentially its newly observed condensate-forming behavior may influence the oligomerization and structural remodeling of HTT1a (Fig. 8).

**Figure 8.**
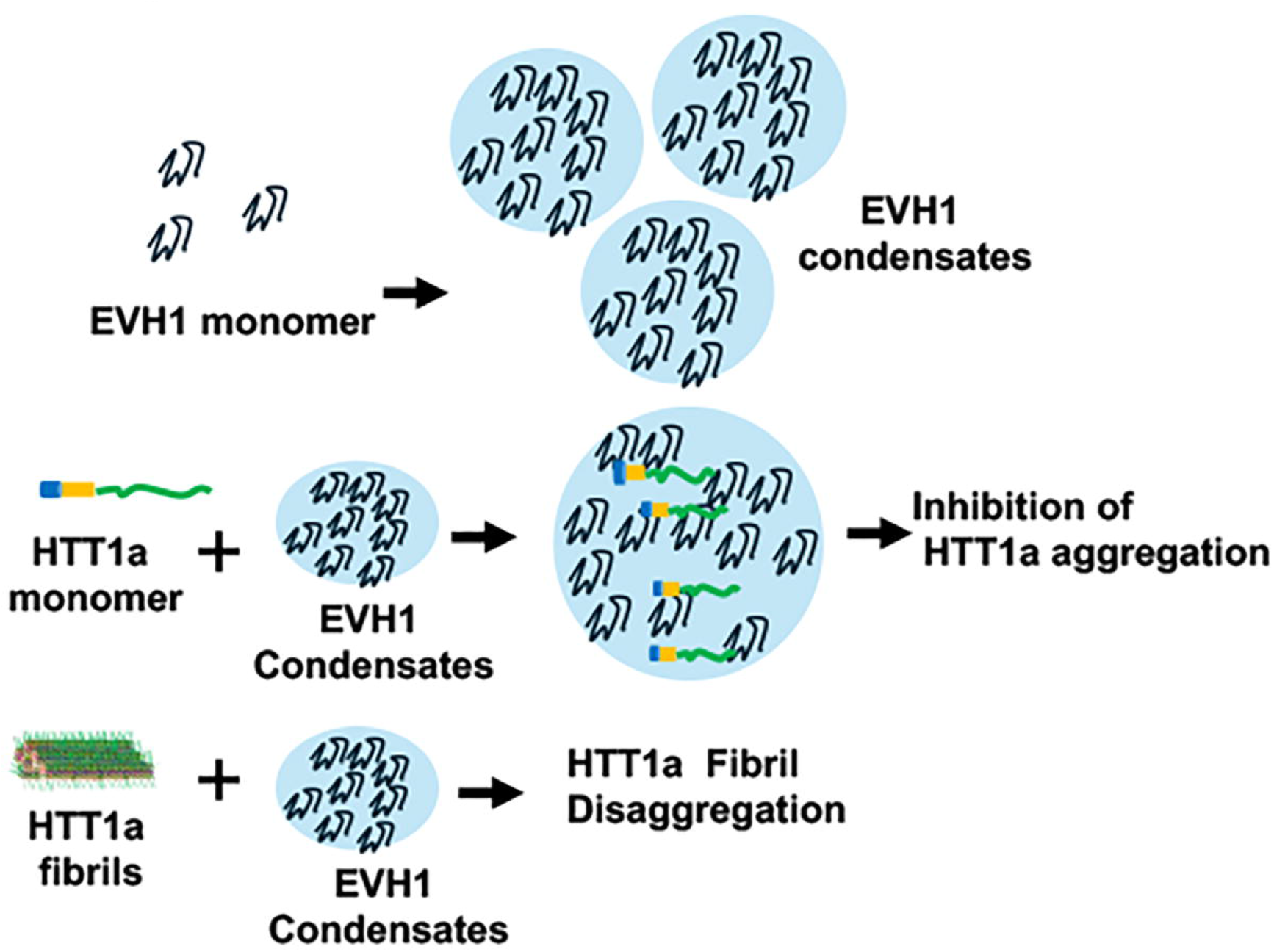
Schematic representation of ENAH-HTT1a interactions. Proposed model in which EVH1 condensate-like assemblies suppress HTT1a aggregation. EVH1 forms condensate-like compartments that recruit soluble HTT1a monomers through PRD-dependent interactions, limiting their availability for nucleation and fibril growth. EVH1 also binds HTT1a fibrils and may inhibit further elongation or promote fibril remodeling, shifting HTT1a from aggregation-prone assemblies toward EVH1-bound, non-fibrillar states.

Mass spectrometry of His-PRD pulldowns identified ENAH as a significant interactor across multiple neuronal fractions and as the top candidate in membrane fractions, alongside VASP and established PRD-binding proteins such as profilin (Supplementary Figure 1;10). These newly discovered binders along with profilin reinforce the concept that the PRD serves as a domain for mediating interactions with multiple polyP-binding proteins (*9*). ENAH and VASP share conserved EVH1 domains that recognize polyP motifs. Given the repetitive and structurally accessible nature of the PRD in HTT1a, particularly in fibrillar assemblies (*21, 23*), EVH1 may engage this region with high avidity through multivalent interactions. The enrichment of ENAH in the PRD pulldowns from multiple organelles suggests that HTT1a may more broadly interact with cytoskeletal and membrane-associated regulatory networks than previously appreciated. In dot blot assays, recombinant EVH1 displayed stronger signal with HTT1a fibrils over non-assembled species. This could arise from multivalent engagement of repetitively displayed PRD motifs on the fibril surface, where the PRD remains accessible, as previously demonstrated by the binding of polyproline-directed antibodies to HTT1a assemblies (*23*).

An unexpected finding was the intrinsic condensation behavior of EVH1 under macromolecular crowding conditions. In the presence of PEG, EVH1 rapidly formed condensate-like assemblies that recruited soluble HTT1a monomers. Although phase separation has been described for related cytoskeletal regulators such as VASP, condensation of ENAH has not been previously reported (*24*). Recruitment of HTT1a into EVH1 condensates suggests that EVH1 may act as a scaffold, concentrating PRD-containing substrates within dynamic compartments. Such compartmentalization could influence HTT1a sequestration, remodeling, or coupling to cytoskeletal and trafficking pathways. In the context of emerging evidence that biomolecular condensates contribute to protein quality control, EVH1-mediated compartmentalization may represent an endogenous homeostatic mechanism against mutant HTT accumulation.

Consistent with its PRD-binding activity, EVH1 significantly inhibited HTT1a-Q46 aggregation in Thioflavin T assays and prevented fibril formation as assessed by TEM. Importantly, EVH1 itself did not aggregate, indicating specificity of the inhibitory effect. Moreover, EVH1 promoted rapid unbundling and remodeling of preformed HTT1a fibrils, with structural disruption evident within minutes and progressing over extended incubation. These findings suggest that EVH1–PRD interactions destabilize fibrillar architecture, potentially by perturbing PRD-mediated intermolecular packing. Multivalent engagement of exposed PRD regions may shift the equilibrium toward non-fibrillar or more dispersed states.

In human stem cell–derived neuronal progenitors expressing HTT1a-72Q, ENAH robustly colocalized with HTT1a in the cytoplasm and neuritic extensions. This spatial overlap indicates that the interaction observed biochemically also occurs in a relevant neuronal HTT1a model. ENAH enrichment in HTT1a-positive regions suggests preferential association with assembly-prone or assembled species rather than diffuse monomers. Given the dynamic localization of actin-regulatory proteins, ENAH recruitment to HTT1a assemblies may reflect a novel neuronal response to aberrant protein accumulation, which remains to be investigated in future studies.

Collectively, these findings identify ENAH as a previously unrecognized regulator of HTT1a assembly dynamics and expand the functional repertoire of EVH1 domains beyond classical actin-associated signaling. Given the accessibility and repetitive display of the PRD in HTT1a fibrils, this region represents a strategic node for modulating HTT1a proteostasis. Future studies should determine whether full-length ENAH recapitulates the remodeling activity observed with isolated EVH1, how EVH1 condensation behavior is regulated in neurons, and whether ENAH modulates HTT1a toxicity in vivo. Our findings position ENAH and related EVH1-containing proteins as endogenous modulators of HTT1a aggregation and suggest that future investigations may identify potential targets for limiting HTT1a oligomerization and neurotoxicity.

### Author contributions

Author contributions: JV, AR, PP, and BQ performed the experiments. JV, TC, RL, and AK designed the experiments and analyzed the data. JV, RL, and AK wrote the manuscript.

## Acknowledgements

We thank Dr. Amy Keating (MIT) for providing the EVH1 bacterial expression vector. This work was supported by an NIH R01 grant to AK and RL.

